# RNAdvisor 2: A unified platform for RNA 3D model quality assessment using metrics, scoring functions, and meta-metrics

**DOI:** 10.1101/2025.09.01.673495

**Authors:** Clément Bernard, Guillaume Postic, Sahar Ghannay, Fariza Tahi

## Abstract

RNA is a molecule that performs critical roles in cellular biology, with its function closely dependent on its three-dimensional conformation. Predicting and evaluating RNA 3D structures remains a significant challenge in the field. Although many metrics and scoring functions have been developed to assess structural quality, each offers a different perspective, and no single method has emerged as a definitive standard. To address this, we previously introduced RNAdvisor, a comprehensive and automated software platform to evaluate 3D RNA structures using a wide range of existing quality metrics and scoring functions. This work presents RNAdvisor2, an extended and improved version of the tool. RNAdvisor 2 introduces a web server designed to enhance accessibility and usability for the broader research community. This release includes new scoring functions and integrates the novel concepts of meta-metrics and meta-scoring functions, which unify diverse evaluation criteria into more robust indicators of RNA structure quality. Additionally, the command-line tool has been improved and optimized for greater stability and extensibility, supporting scalable and maintainable future development. The web server, tool and its source code are freely available on the EvryRNA platform: https://evryrna.ibisc.univ-evry.fr.

## Introduction

The accurate evaluation of RNA three-dimensional (3D) structures is critical for understanding RNA function and guiding structure prediction methods. As experimental determination of RNA 3D structures remains laborious and limited in throughput, computational predictions have become essential (1). Their evaluation must be assessed with robust, interpretable, and scalable evaluation tools. Over the past decade, a diverse number of evaluation metrics (2–5) and scoring functions (6–9) have been developed to estimate the nativeness and quality of RNA structural models.

Metrics are quantitative measures that assess specific aspects of structural quality by comparing with a native conformation, such as the overall folding (2) or atomic interactions (3, 5). In contrast, scoring functions are reference-free computational methods designed to evaluate and rank candidate models during structure prediction, often by computing key RNA features that relate to structural stability (6, 7, 10, 11). While individual metrics and scoring functions can perform well for specific use cases, their effectiveness and generalizability often vary across datasets and structural contexts. Moreover, no single metric or scoring function can comprehensively capture all aspects of structural quality. This lack of a universal standard leads to a fragmented evaluation landscape, where multiple measures may offer complementary or even redundant insights. Recognising this complexity, recent initiatives such as CASP15 (12) and CASP16 (13) have begun exploring integrated “meta” evaluation strategies, which aim to synthesise diverse quality indicators into unified scores for a unique assessment.

In our previous work (14), we introduced RNAdvisor, a unified command-line tool that gathers in one interface the computation of eleven widely used evaluation metrics and four scoring functions. To support broader accessibility and ease of use, we have also developed a web-based interface for RNAdvisor. This platform allows users to interactively compute and compare a wide range of evaluation metrics and scoring functions. This updated release incorporates two additional evaluation metrics: CLASH-score (15) and LCS-TA (16) as well as eight newly developed scoring functions and their variants, including cgRNASP (9), LociPARSE (8), RNA-BRiQ (17), PAMNet (18), RNA3DCNN (19), TB-MCQ (20), ARES (11), and 3dRNAScore (10). Furthermore, we have restructured RNAdvisor’s architecture to allow independent installation and execution of each metric and scoring function. This modularity enhances usability and flexibility, enabling users to focus on specific methods relevant to their application. A major new feature of RNAdvisor is the introduction of a meta-metric framework, which aims to aggregate multiple evaluation scores into a single interpretable value. We further propose a novel meta-scoring function that ranks RNA structure decoys without requiring a native reference, extending the utility of scoring functions in real-world prediction scenarios. These meta-evaluation strategies represent an important step toward developing consensus-based quality measures that are robust across datasets and modelling approaches. Users can easily customise the “meta” metrics and scoring functions with the help of our integrated and complete tool.

### RNA quality assessment

#### Metrics and meta-metrics

To assess whether a predicted RNA tertiary structure is close to its native fold, multiple metrics have been developed. They can be general, telling how well the prediction falls into the global conformation (RMSD, *ϵ*RMSD (21) and CLASH-score (15)). Other metrics, inspired by protein metrics, consider the alignment of structures to evaluate a predicted structure (TM-score (2), GDT-TS (22), CAD-score (4) or LDDT (5)). Nevertheless, proteins and RNAs have differences that limit the adaptation of protein metrics to RNAs. One of the significant difference lies in folding stabilisation, where RNA structure is maintained by base pairing and base stacking, compared to hydrogen interactions for protein structures. Therefore, metrics have been developed to fit the RNA specificities, considering the different types of interactions (INF (3), DI (3)) or torsional angles (MCQ (23), LCS-TA (16)). A more complete description of the metrics is available in our previous work (14). In this new version of RNAdvisor, we have added the LCS-TA (with the possibility to choose the threshold of continuity), the CLASH-score and the C-LDDT (coarse-grained LDDT). We also changed the library used for the TM-score (from OpenStructure (24) to US-Align (25)). Each metric represents the result of years of development, but no single value can appropriately assess RNA structural quality. While some metrics may be redundant, others can offer complementary insights. Choosing the appropriate metric is challenging and highly dependent on the specific context. To better leverage their strengths, we can consider designing meta-metrics that combine the benefits of existing approaches. The goal is to have a unique value that would give an overview of the quality of a structure, and thus be used to assess the quality of a prediction. In order to aggregate and consider the variability of the metrics, the Z-score can be used to normalise them. The Z-score is a statistical measure that describes the number of standard deviations a data point is from the mean. Given a metric *M*, the Z-score is defined as:

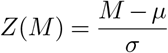

with *µ* the mean and *σ* the standard deviation of the metric. In CASP 15 (12), a meta-metric was proposed using a weighted sum of Z-scores of different metrics:

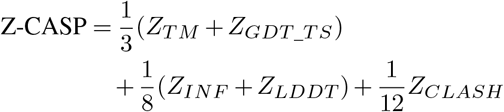

In CASP16 (13), another weighted sum of Z-scores was proposed, but with different weights:

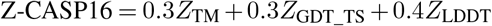

The weights of each meta-metric are defined based on the importance of the metric in assessing the quality of a structure, and are often chosen based on expert knowledge or empirical results. In CASP15 (12), the weights were chosen based on the assumption that highly accurate results were not expected. The weights favoured metrics capturing the global fold (higher weights for TM-score and GDT_TS) compared to those assessing local details.

Instead of selecting specific weights, another solution is to sum all available metrics, hoping that the overall sum provides a good overview of structure quality. Therefore, we propose to use a weighted sum of Z-scores from different metrics, defined as:

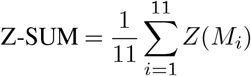

The considered metrics are RMSD, *ϵ*RMSD, CLASH-score, TM-score, GDT-TS, CAD-score, LDDT, P-VALUE, INF, MCQ, and LCS-TA. Some metrics require parameter choices, such as the threshold for LCS-TA or the choice of coarse-grained atom for TM-score. For LCS-TA, we used a 10^*°*^ threshold for angle conservation to create a stricter metric. We used the default 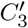 for the TM-score.

Instead of using Z-scores, we can also apply min-max normalisation to each metric:

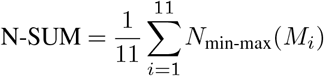

where *M*_*i*_ is the *i*-th metric, and *N*_min-max_ is the normalisation function that scales the metric between 0 and 1. For descending metrics (where lower values indicate better performance), the values are reversed so that higher normalised values always indicate better predictions. The goal is thus to maximise the value of the meta-metric, which reflects prediction quality. This normalisation is intuitive, since all metrics are scaled between 0 and 1.

The advantage of Z-score normalisation is that it accounts for the distribution of each metric, offering a more nuanced assessment of structure quality. However, Z-scores may include negative values and are harder to interpret. Min-max normalisation is easier to understand but can be biased by outliers. Each method has trade-offs: Z-scores reflect variability, while min-max normalisation offers intuitive comparability. This is why we propose to keep both of these normalisations and use them differently depending on the context.

As seen with the weight change between CASP15 and CASP16, the choice of weights in meta-metrics can significantly affect the structure rankings. This choice is not trivial and depends on the dataset. Thanks to our RNAdvisor 2 wrapper, which supports different metrics, it becomes easy to use different meta-metrics that can be adjusted on the study case.

#### coring functions and meta-scoring functions

Dissimilarity metrics can compute the nativity of RNA molecules, but require a known solved reference structure. This requirement is challenging as the number of solved structures is low. Furthermore, computational methods usually predict multiple conformations that need to be ranked (26–29). Adapting the structure’s free energy has become a standard in structures’ ranking, filtering, and confidence assessment. These predictive quality measurements are either knowledge-based approaches (3dRNAScore (10), *ϵ*SCORE (21), RASP (7), DFIRE-RNA (30), rsRNASP (6), cgRNASP (9)) that rely on statistical potentials or deep learning ones (RNA3DCNN (19), ARES (11), PAMNet (18), TB-MCQ (20), LociPARSE (8)). Compared to our previous work, we have added cgRNASP, RNA3DCNN, ARES, PAMNet, TBMCQ and LociPARSE. Details on the description of each scoring function are available in our previous work (14). The added scoring functions are described in the Supplementary file.

Similarly to the meta-metrics, we can define a meta-scoring function that combines the different scoring functions. To the best of our knowledge, we have not seen any contribution using this principle for the scoring functions. We can use the same idea for the meta-metrics by using a weighted sum of the Z-scores of the different scoring functions. The choice of the scoring function is not trivial, as it can change the ranking of the structures. We can also use a min-max normalisation of the scoring functions, which would give a better overview of the quality of the structure. Therefore, we have created a meta-scoring function based on four existing scoring functions: three of them based on the highest correlation with the sum of metrics and the last one with the highest correlation with the angle-specific MCQ metric. More precisely, our meta-scoring function, named Z-META, is composed of:

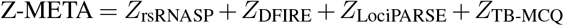

with *Z*_rsRNASP_ with the Z-score for the rsRNASP scoring function (respectively for the other scoring functions). We also extend this formulation with the min-max normalisation, which we call N-META.

### RNAdvisor 2 tool

Our previous version of our tool, RNAdvisor (14), enabled the computation of state-of-the-art metrics and scoring functions in one command line. It grouped eleven metrics and four scoring functions. In this new version, we have increased the number of available metrics to thirteen metrics and eighteen scoring functions (counting the variant tools available for each scoring function).

While the previous version of RNAdvisor used one Docker container (31) where all the metrics were available under one image, the new version provides a unique image for each available metric and scoring function. The architecture is shown in Figure 1. It allows users to study a specific metric or scoring function if needed, and makes the extension much easier to add new metrics or scoring functions. All the Docker images have a similar Python wrapper inside to enable easy usage. Users are not required to have a unique image with all the metrics available to compute one unique metric: specific Docker containers are downloaded when the user requires them. The wrapper uses Python and is easily installable. All laborious installations are already done, and the volume mapping between local folders and Docker containers is done in the Python wrapper. The code can be run in parallel, and each container’s size is kept as small as possible to minimise computation times.

**Fig. 1.**
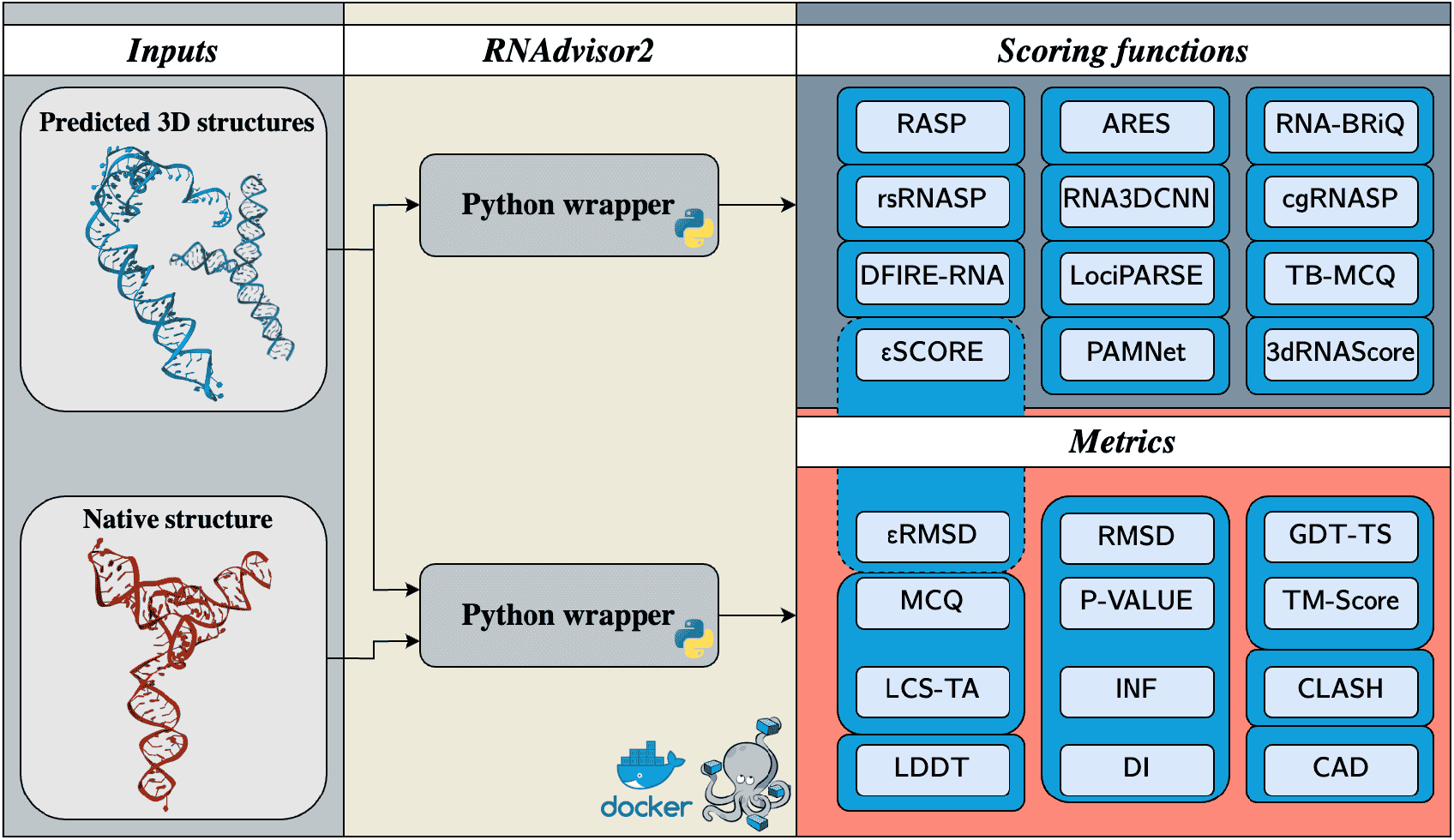
Schema of RNAdvisor 2. A wrapper code that gathers libraries for the assessment of RNA 3D structures in one interface. A Python wrapper is used, easily installable with pip. Each scoring function and metric is in its own Docker container, which allows the user to choose which metric or scoring function to use.

In addition to the command-line tool, we provide a web server (Figure 2) for the computation of metrics and scoring functions, accessible with the source code in the EvryRNA platform https://evryrna.ibisc.univ-evry.fr/evryrna/. The web interface offers a user-friendly alternative for individuals less familiar with command-line operations. It is particularly well-suited for evaluating a small number of RNA 3D structures. The user can choose between two use cases: metrics or scoring functions (Figure 2 A). Then, a list of metrics or scoring functions is available. The user can then upload their RNA 3D structures. In the case of the metrics scenario, the server’s output is a dataframe with the different metrics, followed by a barplot of the N-SUM meta metric (better for visualisation than the Z-SUM) (Figure 2 B). The Z-score and normalised metrics are directly available in the dataframe: the user can then download and create their own meta-metric easily. A polar plot is also provided with INF submetrics (INF_*W C*_, INF_*nW C*_ and INF_*ST ACK*_) (Figure 2 C)). A time plot also shows the time spent on each metric. In the scoring function scenario, the dataframe is shown as the first output and a barplot with normalised scoring functions (by the min-max values over the set of structures provided). If the TB-MCQ is selected, a specific figure is shown with a polar plot for the different torsional angles Figure 2 D) and the TB-MCQ per position (Figure 2 E). Not all metrics and scoring functions are provided in the web server. We only considered the fastest ones, while the rest are available using the command line.

**Fig. 2.**
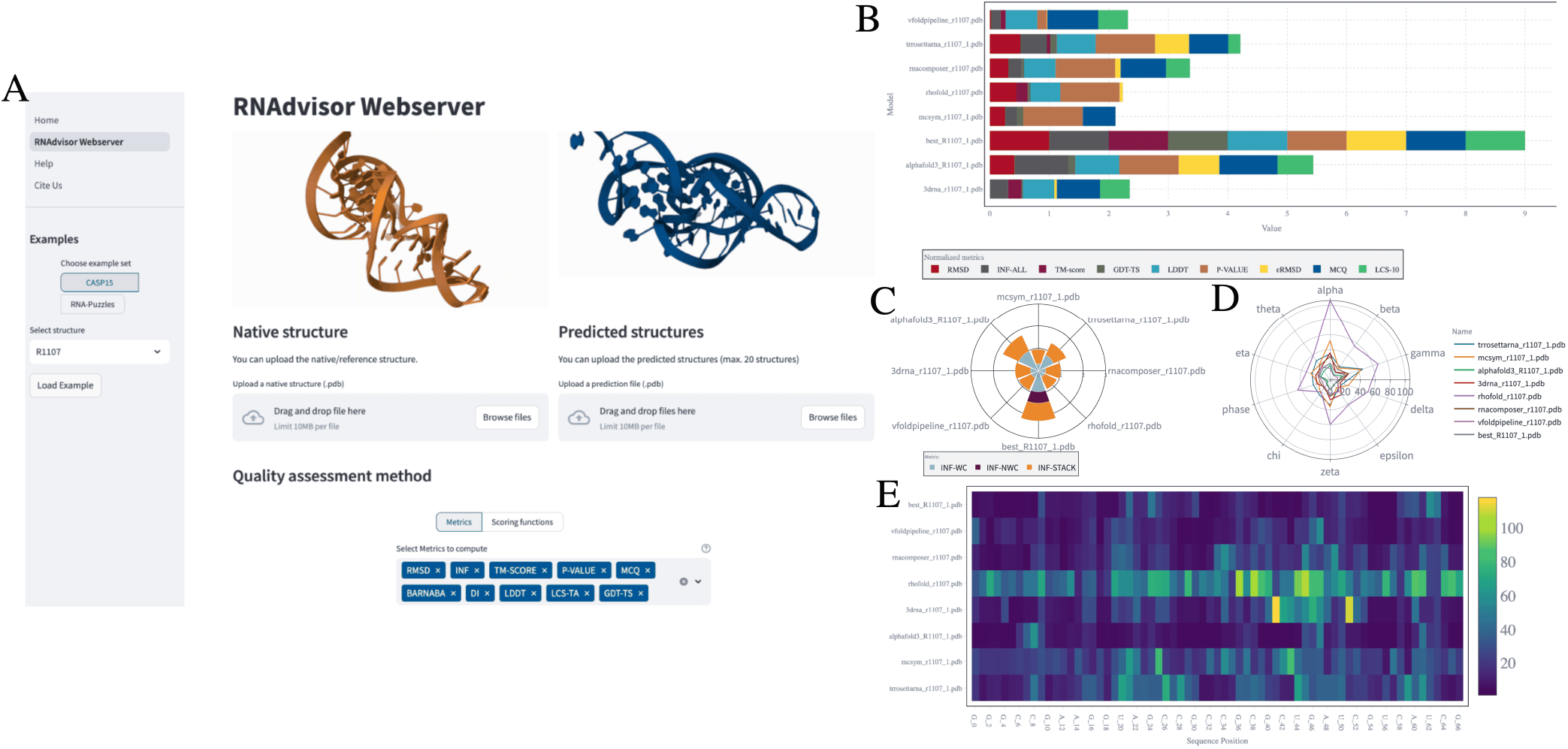
Screenshot of RNAdvisor 2 web server for challenge R1107 of CASP-RNA (32). Left: Inputs. Right: Outputs. A) Main page where the user can add inputs or load available examples. B) Bar plot with the sum of normalised metrics when running the metrics computation. C) Polar plot with INF metrics for the metrics computation. D) TB-MCQ (20) for each torsional angle for the scoring function scenario. E) TB-MCQ (20) per position when the user chooses to compute the TB-MCQ for the scoring function scenario.

### Benchmark

This section provides a complete and exhaustive benchmark of the scoring functions and metrics. We compare existing metrics and provide a benchmark of the scoring functions by comparing their performance on different datasets, including our meta-scoring function. In the literature (6, 8, 11, 19), the evaluation of scoring functions is often done by comparing the rank of the native structure in a set of decoys. This evaluation can be strengthened by adding the correlation to existing metrics or comparing the top-ranked decoys with the native structure.

To evaluate the link between metrics and scoring functions, we compared the PCC (Pearson correlation coefficient) and ES (enrichment score) between each computed metric. More details on their formulation are provided in the previous paper (14). The PCC is used to assess the correlation, whereas the ES counts for the number of common 10% structures based on the scoring function or metric ranking.

We used four datasets, Test Set I to Test Set IV, to assess the relations between scoring functions and metrics. The first two datasets have decoys generated by two strategies widely used to compare scoring functions (6, 7, 11, 19). The last two datasets are a real-case scenario where 3D structures from different model predictions should be ranked by nativity. Compared to our previous work (14), we added one dataset, Test Set IV, to our benchmark. The description of the first three datasets can be found in RNAdvisor(14).

**Test Set IV**^1^ is the CASP15 dataset. It is composed of 12 RNAs with decoys generated by different teams participating in the CASP15 competition. There is a higher number of decoys for each RNA compared to Test Set III (more than a hundred per target), and the difficulty of the prediction is higher.

In this section, we first present the link between the different metrics. Then, we discuss the quality of the scoring function using the ranking of the native structure and its correlation with existing metrics. Finally, we discuss the computation time for the different metrics and scoring functions.

#### Link between metrics

To assess the quality of RNA structures given a reference, metrics consider different specificities like base interactions, angle conservation, distance deviation, etc. We provide the mean of the PCC and ES averaged over the four test sets in Figure 3, which is an updated version of our previous work (14) (one more dataset and three more metrics). The results for each dataset are provided in Figure S1, S2, S3 and S4 of the Supplementary file. We discuss here the new results obtained. The added results are the correlations of C-LDDT, LCS-TA (with a strict threshold of 10°, which we refer to as LCS-10), P-VALUE and CLASH-score. Surprisingly, the C-LDDT is not as correlated with its full version (LDDT) as expected. It has the highest correlation with GDT-TS, CAD-score and *ϵ*RMSD (PCC of 0.84, 0.86 and 0.85, respectively). LCS-10 has low correlation with RMSD, LDDT and INF_*all*_ but is more correlated to MCQ (PCC of 0.72 and ES of 5.34) and TM-score (PCC of 0.7 and ES of 5.01). Its link to MCQ is expected, while its correlation with TM-score is more surprising. It might be explained by the fact that a long sequence of conserved angles might be linked to backbone conservation. Since the P-VALUE measures the probability that a given structure performs better than expected by chance, values close to zero indicate good predictions. In contrast, higher values suggest the prediction is no better than random. Therefore, the P-VALUE does not serve well for selecting highly discriminative decoys but acts as a baseline criterion for identifying acceptable predictions. The CLASH-score is not correlated to the other metrics, as it is considered more as a quality metric without a reference structure. Even native structures can have small clashes, which makes it less relevant to compare with other metrics. It is worth noting that the CLASH-score correlates with the LDDT (PCC of 0.70), but fails in terms of ES (ES of 2.08).

**Fig. 3.**
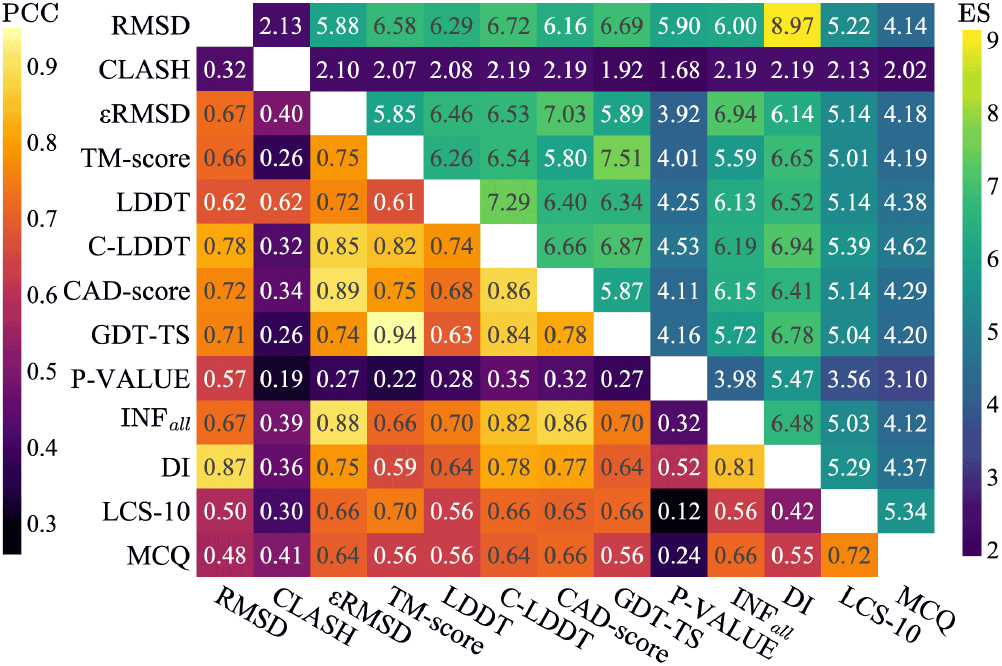
ES and PCC scores for each metric averaged over the four test datasets (TSI, TSII, TSIII and TSIV). The lower half of the matrix represents the PCC, while the upper half corresponds to the ES score. The diagonal has a PCC of 1 and ES of 10.

The previous results are still valid (with the addition of one dataset). We discuss the link between the different meta-metrics in the Supplementary file.

The correlation between the different metrics is not perfect (no ES of 10 or PCC of 1), meaning that every metric can help assess predicted model quality. The torsional-based metrics tend to be less correlated with the other metrics, which is not surprising, as they do not use the same structural features for comparison. The angle conservation is not mainly included in other metrics computation, which could explain this behaviour. The CLASH-score is a quality assessment that is more like a condition to be respected for a structure rather than an absolute metric to rank them.

#### Scoring function ranking

The aptitude of a scoring function to classify native and near-native structures is essential for developing models. Table 1 shows the number of times the native structures are found with the best scoring function value for each decoy (it can either be the lowest or the highest value, depending on the scoring function).

**Table 1.** Number of native structures found with the best scoring function value for each dataset. It corresponds to the number of times the native structure has the lowest-scoring function value among the decoys. Methods are ranked by the overall sum. The number of structures is 85 for Test Set I (TSI), 20 for Test Set II (TSII), 21 for Test Set III (TSIII) and 12 for Test Set IV (TSIV). The total is out of 138 structures. Scoring functions are either knowledge-based (KB) or deep learning (DL).

|  | TSI | TSII | TSIII | TSIV | Total |
| --- | --- | --- | --- | --- | --- |
|  | 85 | 20 | 21 | 12 | 138 |
| rsRNASP (6) [KB] | 77 | **16** | **18** | 0 | **111** |
| εSCORE (21) [KB] | **85** | 9 | 5 | **4** | 103 |
| DFIRE (30) [KB] | 81 | 10 | 10 | 0 | 101 |
| Z-META [KB/DL] | 67 | 13 | 12 | 0 | 92 |
| RASP (7) [KB] | 83 | 2 | 2 | 0 | 87 |
| N-META [KB] | 61 | 12 | 12 | 0 | 85 |
| PAMNet (18) [DL] | 70 | 6 | 5 | 3 | 84 |
| cgRNASP (9) [KB] | 34 | 15 | **18** | 1 | 68 |
| cgRNASP-PC (9) [KB] | 32 | 11 | 16 | 1 | 60 |
| RNA-BRiQ (17) [KB] | 39 | 7 | 8 | 0 | 54 |
| 3dRNAScore (10) [KB] | 52 | 0 | 0 | 0 | 52 |
| RNA3DCNN_MD (19) [KB] | 35 | 4 | 6 | 0 | 45 |
| LociPARSE (8) [DL] | 18 | 14 | 8 | 2 | 42 |
| RNA3DCNN_MDMC(19)[KB] | 24 | 5 | 11 | 0 | 40 |
| TB-MCQ (20) [DL] | 28 | 5 | 3 | 1 | 37 |
| cgRNASP-C (9) [KB] | 21 | 6 | 9 | 1 | 37 |
| ARES (11) [DL] | 12 | 1 | 0 | 0 | 13 |

The best method is rsRNASP, which finds 111 out of 138 native structures. It has the best results for Test Set II and III, but is not the best for Test Set I nor Test Set IV. Test Set I is the one with the most near-native decoys, whereas Test Set IV is the one with the longest RNAs. Incorporating statistical potentials that weigh differently in short, mid-range, and long interactions, like rsRNASP, may not be the best choice for very close decoys. Its light version, cgRNASP, performs as well as rsRNASP for Test Set III (with 18 out of 21 native structures), but fails for Test Set I (with 34 out of 85 native structures). The best method for Test Set IV is *ϵ*SCORE, which finds 4 out of 12 native structures. This is still less than 50% of the native structures of the dataset, which is overall low. Our Z-META and N-META methods are ranked fourth and sixth, respectively, with 85 and 92 native structures identified. This outcome can be attributed to the fact that these methods rely on a combination of existing scoring functions. As a result, their performance may be weakened by the averaging process, which can dilute the influence of the most effective individual scores. The best deep learning scoring function is PAMNet, which finds 84 out of 138 native structures. It performs well on Test Set I but fails on Test Set II and III. It is the second-best for Test Set IV. Our scoring function, TB-MCQ, is not the best for ranking native structures, as it only considers the torsional information. The worst scoring function in this case is ARES, which finds only 13 out of 138 native structures. This could be explained by the fact that it has memorised the type of decoys from its training set, which differs from these datasets. Also, the range of values of ARES is quite narrow, leading to few values to rank the structures.

The results on the evaluation of the rank of the native structures for the scoring functions show that the best scoring functions are not deep learning methods, but rather statistical potentials. Even the use of a meta-scoring function does not improve the results. Indeed, our meta-scoring function involves two deep learning approaches that perform poorly in ranking the native structure. The best deep learning method is PAMNet, which ranks 7th out of the scoring functions. It does not mean that the scoring function is not relevant, but more than the assessment of the native structure is harder for them, as it requires having a fine assessment of this specific structure. Results highly depend on the nature of the decoys for the specific dataset: a scoring function can perform very well for a dataset, but poorly on another one.

We also observe the ranking quality for the different scoring functions in the Supplementary file.

#### Scoring functions and metrics relationship

Comparing the scoring functions based only on the rank of the native structure is not enough to assess their quality. Indeed, the evaluation is done only based on the native structure, which is hard to achieve. Another way to assess the quality of scoring functions is to compare them with existing metrics, using PCC or ES with the metrics. We computed the ES and PCC scores for each data set and each available scoring function and metric. We considered for the metrics the RMSD, INF (also named INF_*all*_, as we considered the averaged value over base-pairing and base-stacking interactions), DI, MCQ, TM-score, GDT-TS, CAD-score, LDDT, C-LDDT, CLASH-score and *ϵ*RMSD. We have also added the meta-metric Z-SUM, which serves as the final value of ranking across all scoring functions. The ES and PCC scores averaged over the three datasets are shown in Figure 4. The results for each dataset are provided in Figure S9, S10, S11 and S12 of the Supplementary file.

**Fig. 4.**
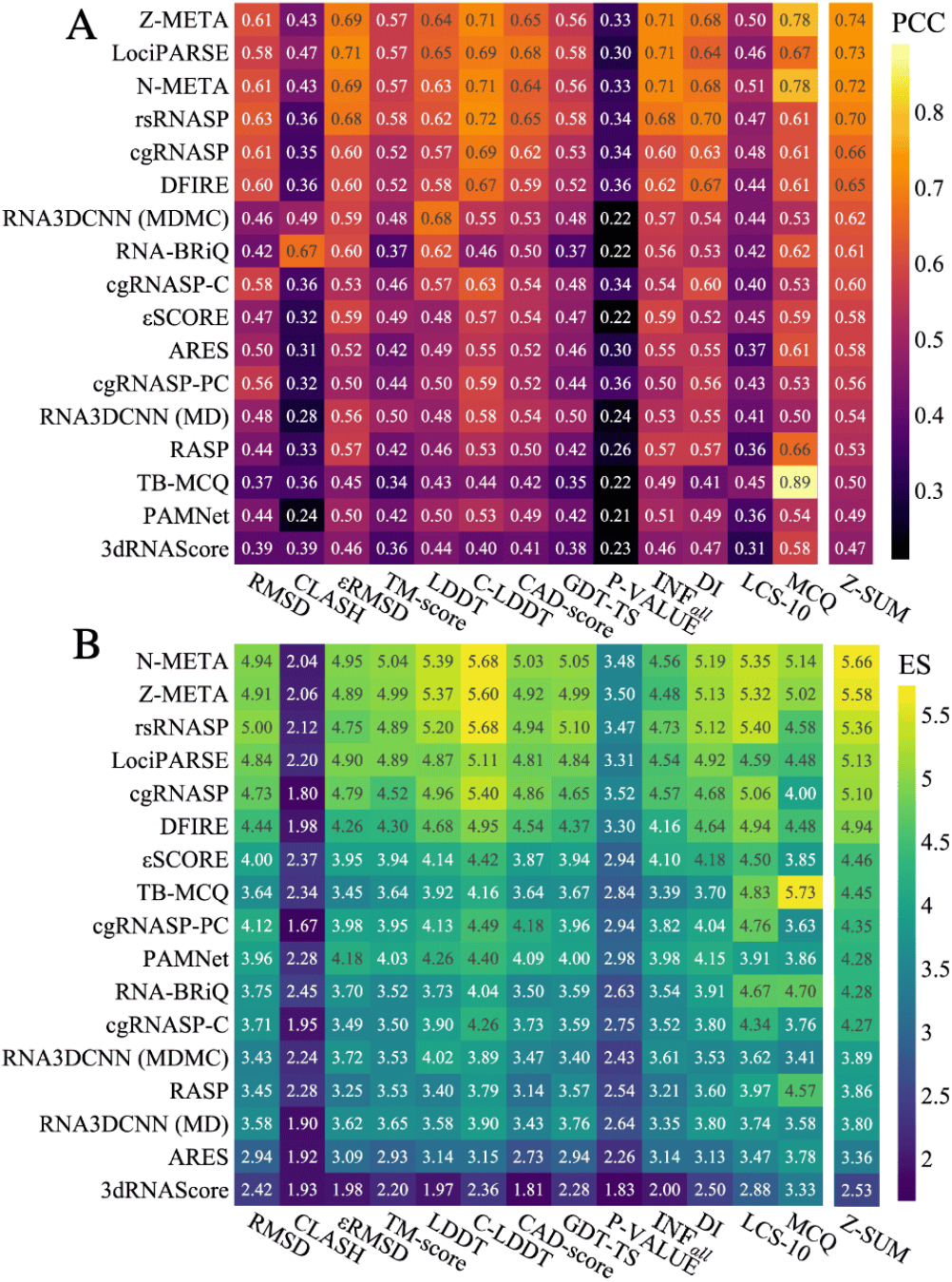
Link between the scoring functions and the metrics averaged over the four datasets. A) PCC and B) ES scores. Scoring functions are sorted by the Z-SUM (last column). Lighter colors indicate stronger link with the metrics, highlighting better overall performance.

The two meta-scoring functions, N-META and Z-META, consistently emerge as top performers in correlation and enrichment. They exhibit the highest PCCs across nearly all structural quality metrics, with values exceeding 0.70 of PCC (and ES higher than 5) for INF_*all*_, C-LDDT and MCQ. They also show the highest correlation (ES and PCC) for the meta metric Z-SUM. This strong alignment highlights their robustness and ability to generalise to the different decoys of the datasets. Among individual methods, LociPARSE and rsRNASP stand out for their excellent balance between correlation and enrichment. They achieve high PCCs (up to 0.70 with DI and 0.72 with C-LDDT) and maintain some of the highest ES values observed across several core metrics (RMSD, INF_*all*_, GDT-TS, C-LDDT and LCS-10). These results position rsRNASP and LociPARSE as high-quality scoring functions supporting robust structure quality assessment. Their approach is very different: rsRNASP is a statistical potential while LociPARSE is a deep learning method. DFIRE and cgRNASP show slightly lower but still competitive results. Both maintain PCCs in the 0.6–0.7 range for several metrics and ES values around 4.5–5.4, suggesting their general reliability. In contrast, deep learning-based methods such as RNA3DCNN (MD/MDMC) and PAMNet occupy a mid-range position in both analyses. While they exhibit moderate PCCs (generally 0.50–0.60), their ES values hover around 3.5–4.0. RNA3DCNN (MDMC) performs relatively better for LDDT in terms of PCC, suggesting it may be particularly attuned to local structural features, but lacks broader generalisation across all metrics. A more nuanced case is TB-MCQ, which, despite moderate PCCs across most metrics, achieves an exceptionally high ES (5.73) and PCC (0.89) for MCQ. This indicates that while TB-MCQ is highly specialised and effective for MCQ-based ranking, it does not generalise well to other structural assessment criteria. At the lower end of the performance spectrum, 3dRNAScore, ARES, and RNA-BRiQ display consistently low PCCs and ES values. 3dRNAScore, in particular, scores below 0.50 in nearly all correlations and below 3.3 in ES, suggesting limited applicability in rigorous structural evaluation. These results suggest these methods are suboptimal for correlation-based analysis and enrichment-based structure ranking. We also observe very low correlation for P-VALUE and CLASH-score among all the scoring functions. It can be explained by the P-VALUE being only values between 0 for non-random predictions and 1 for random ones, which is not what is aimed to provide with scoring functions. The CLASH-score is also not what tends to be reproduced with scoring functions, as it is more a condition to have rather than a criterion to discriminate structures.

#### Computation time

Models for predicting 3D structures are usually slow, and even slower when the sequence length increases (1). The computation time of the scoring functions, used to rank predictive candidates, should not be a bottleneck for selecting decoys for created models.

We tracked the inference computation time for each scoring function for RNAs of different lengths. We took as a benchmark the chain A of RNA 3f1hA (size 2878 nucleotides) from Test Set I. We randomly created 20 substructures for each step of 100 nucleotides from 100 to 2500. We tracked and averaged the time required to compute the scoring functions and metrics (each substructure having a different complexity, we tracked the mean and standard deviation). It leads to Figure 5. It shows the low computation time for metrics like *ϵ*RMSD, GDT-TS, MCQ, DI, RMSD, and INF. The LCS-TA, CLASH-score, CAD-score and LDDT have a higher computation time than the other metrics. LCS-TA has a computation of around 8 minutes for a sequence of 2500 nucleotides, and its computation time depends on the quality of the structure. CLASH-score has a computation time of around 1 min for a sequence of 2500 nucleotides, while CAD-score and LDDT have computation times of 2 min and around 3 min, respectively. The choice of implementation is also important. Previously (in the first version of RNAdvisor), we used the implementation of TM-score from the open structure software (5), which turned out to be very slow. We switched to implementing TM-score from the US-align software (25), which is much faster.

**Fig. 5.**
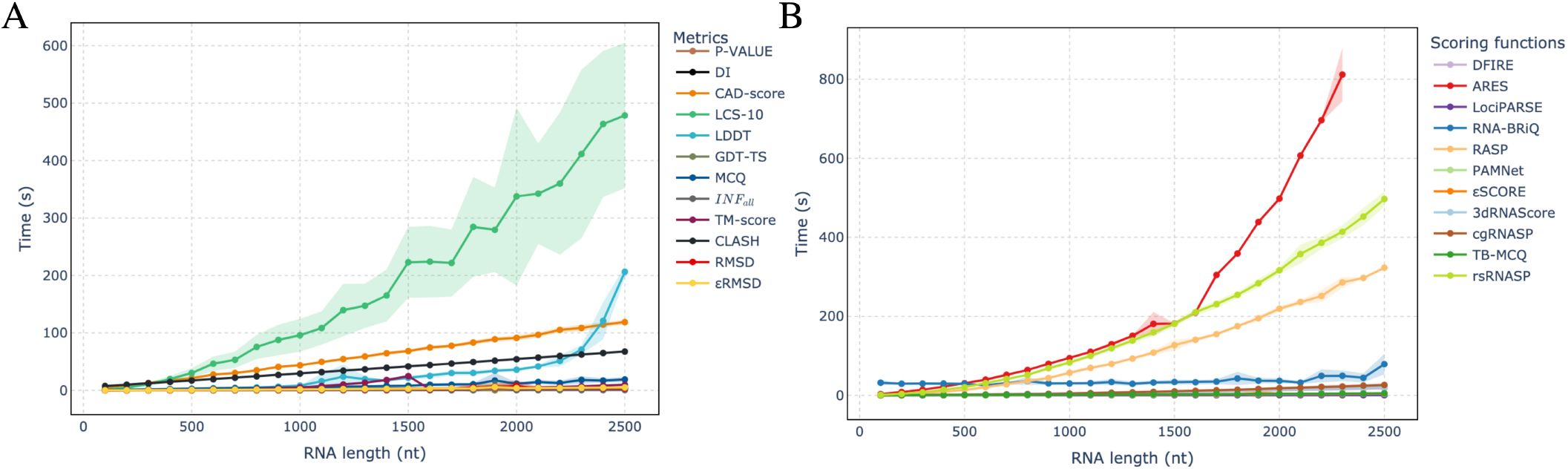
Mean computation time (with its standard deviation) depending on the number of nucleotides in RNA sequences for substructures from RNA 3f1hA (Test Set I). A) Time executions for metrics. B) Time executions for scoring functions.

Regarding the scoring functions, most of them have a low computation time of less than 50 seconds for a sequence of 2500 nucleotides. Nonetheless, three scoring functions have a high computation time: ARES, RASP and rsRNASP. ARES has a computation time of around 13min for a sequence of 2300 nucleotides (we had errors for structures of higher length), while rsRNASP and RASP have computation times of 8min20 and 6min48, respectively. The deep learning scoring functions could be enhanced using GPUs, but we did not get the GPU to work for the inference. This computation time is not scalable for the development of high-resolution models. For instance, if a predicted model generates 1000 structures of 2300 nucleotides and then tries to select the best ones with ARES, it will take more than nine days to compute. This scalability issue is a problem for the use of ARES in the development of large-scale models.

Among the scoring functions, rsRNASP and LociPARSE consistently showed strong performance, offering distinct but complementary advantages. rsRNASP, based on statistical potentials, achieves excellent native structure ranking and robust correlations with key structural quality metrics, but its computation time increases with RNA length. In contrast, LociPARSE, a deep learning-based model, matches rsRNASP’s accuracy while maintaining fast inference times, making it more suitable for large-scale evaluations.

#### Example of use

To demonstrate the interest of our tool for quick benchmarking of RNA 3D structures, we show in Figure 6 the N-SUM with benchmarked models used in previous works(1, 33) for the CASP15 dataset. This meta-metric is easily interpretable, and the cumulative sum enables a quick overview of the leading methods. For instance, in this dataset, we can see that AlphaFold 3 (26) does not outperform the best models from CASP15, nor is Boltz-1 (34) an open-source equivalent.

**Fig. 6.**
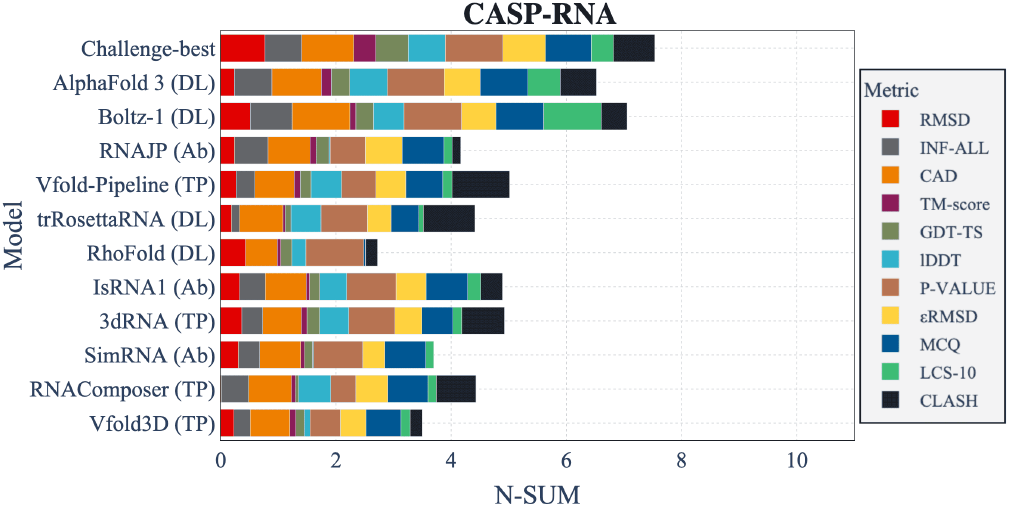
Cumulative normalised metrics for CASP-RNA (32) from benchmarked models used in previous works(1, 35). Each metric is normalised by the maximum value, and the decreased metrics are inverted to have better values close to 1. Challenge-best means the best solutions from the CASP-RNA competition. The types of methods are also mentioned with the abbreviations DL for deep learning, TP for template-based and Ab for ab initio.

## Conclusion

RNAdvisor is a tool designed to consolidate a wide range of existing metrics and scoring functions into a single, user-friendly interface for assessing RNA 3D structural quality. Each metric and scoring function reflects years of research by different scientific groups. By making them accessible through a single command-line interface, RNAdvisor facilitates straightforward use and is particularly well-suited for automating benchmarking workflows. This is especially important given the growing number of RNA 3D structure prediction methods.

In this work, we demonstrate the utility of RNAdvisor 2 through a use case that explores the relationships among various metrics and scoring functions. We also introduce a proof of concept for meta-metrics and meta-scoring functions.

We provide a web-based interface to support users less familiar with command-line tools, making it easier to explore and compare RNA 3D structures. This web server offers a streamlined and intuitive environment for uploading, analysing, and comparing RNA 3D structures. It is particularly helpful for experimental biologists looking for quick and accessible insights without the need for software installation or command-line operations. The web interface replicates the core functionalities of the command-line version and adds visualisation tools to further enhance interpretation and user experience.

Beyond individual scoring functions and metrics, we introduced meta-metrics such as Z-SUM and N-SUM to aggregate multiple quality criteria into a single, interpretable score. These meta-metrics demonstrated strong alignment with traditional evaluation measures, providing a robust and comprehensive framework for structure assessment. There remains a bias in the definition of these meta metrics, which would be the predominance of spatial considerations, as more metrics are based on spatial information. There is also a redundancy in the metrics, which is hardly avoidable, as the metrics are based on different principles.

Similarly, the proposed meta-scoring functions (Z-META, N-META) effectively captured structural quality across diverse decoy sets, with the limitation of direct identification of native structures.

Different combinations of meta-metric and meta-scoring functions can be used, and we did not try different optimised combinations. We decided to use a simple example of meta score to show its interest. If computation time is a bottleneck, one can easily replace rsRNASP with its cgR-NASP equivalent in the meta-scoring function. Also, a better optimised combination of scoring functions can be found. Ideally, to optimise the weights of such meta-scoring functions, another dataset of decoys would be needed to adjust the weights depending on correlation with existing metrics (or meta-metrics). Thanks to our RNAdvisor tool, it becomes very easy to create and custom new meta-metrics and scoring functions.

## Data availability

The RNAdvisor tool, web server and datasets used in this study are available in the EvryRNA platform: https://evryrna.ibisc.univ-evry.fr.

## Footnotes

1 https://predictioncenter.org/casp15/

## Bibliography

1. Clément Bernard, Guillaume Postic, Sahar Ghannay, and Fariza Tahi. State-of-the-RNArt: benchmarking current methods for RNA 3D structure prediction. NAR Genomics and Bioinformatics, 6(2):lqae048, June 2024. doi: 10.1093/nargab/lqae048.

2. Yang Zhang and Jeffrey Skolnick. Scoring function for automated assessment of protein structure template quality. Proteins, 57(4):702–710, December 2004. ISSN 0887-3585. doi: 10.1002/prot.20264.

3. Marc Parisien, José Cruz, Eric Westhof, and François Major. New metrics for comparing and assessing discrepancies between RNA 3D structures and models. RNA (New York, N.Y.), 15:1875–85, 09 2009. doi: 10.1261/rna.1700409.

4. Kliment Olechnovic, Eleonora Kulberkyte, and Ceslovas Venclovas. CAD-score: A new contact area difference-based function for evaluation of protein structural models. Proteins, 81, 01 2013. doi: 10.1002/prot.24172.

5. Valerio Mariani, Marco Biasini, Alessandro Barbato, and Torsten Schwede. lddt: a local superposition-free score for comparing protein structures and models using distance difference tests. Bioinformatics (Oxford, England), 29(21): 2722–2728, 2013. doi: 10.1093/bioinformatics/btt473.

6. Ya-Lan Tan, Xunxun Wang, Ya-Zhou Shi, Wenbing Zhang, and Zhi-Jie Tan. rsRNASP: A residue-separation-based statistical potential for RNA 3D structure evaluation. Biophysical Journal, 121:142–156, 1 2022. ISSN 00063495. doi: 10.1016/j.bpj.2021.11.016.

7. Emidio Capriotti, Tomas Norambuena, Marc A. Marti-Renom, and Francisco Melo. All-atom knowledge-based potential for RNA structure prediction and assessment. Bioinformatics, 27:1086–1093, 4 2011. ISSN 1460-2059. doi: 10.1093/bioinformatics/btr093.

8. S. Tarafder and D. Bhattacharya. lociparse: A locality-aware invariant point attention model for scoring rna 3d structures. Journal of Chemical Information and Modeling, 64(22):8655–8664, Nov 2024. doi: 10.1021/acs.jcim.4c01621.

9. Ya-Lan Tan, Xunxun Wang, Shixiong Yu, Bengong Zhang, and Zhi-Jie Tan. cgrnasp: coarse-grained statistical potentials with residue separation for rna structure evaluation. NAR Genomics and Bioinformatics, 5(1), 03 2023. ISSN 2631-9268. doi: 10.1093/nargab/lqad016.lqad016.

10. Jian Wang, Yunjie Zhao, Chunyan Zhu, and Yi Xiao. 3dRNAscore: a distance and torsion angle dependent evaluation function of 3D RNA structures. Nucleic Acids Research, 43: e63–e63, 5 2015. ISSN 1362-4962. doi: 10.1093/nar/gkv141.

11. Raphael J. L. Townshend, Stephan Eismann, Andrew M. Watkins, Ramya Rangan, Maria Karelina, Rhiju Das, and Ron O. Dror. Geometric deep learning of RNA structure. Science, 373:1047–1051, 8 2021. ISSN 0036-8075. doi: 10.1126/science.abe5650.

12. R Das, RC Kretsch, and AJ Simpkin. Assessment of three-dimensional RNA structure prediction in CASP15. Proteins, 91(12): 1747–1770, 2023. doi: 10.1002/prot.26602.

13. Rachael C. Kretsch, Alissa M. Hummer, Shujun He, Rongqing Yuan, Jing Zhang, Thomas Karagianes, Qian Cong, Andriy Kryshtafovych, and Rhiju Das. Assessment of nucleic acid structure prediction in CASP16. 2025. bioRxiv doi: 10.1101/2025.05.06.652459, 10 May 2025, pre-print: not peer-reviewed.

14. Clement Bernard, Guillaume Postic, Sahar Ghannay, and Fariza Tahi. RNAdvisor: a comprehensive benchmarking tool for the measure and prediction of RNA structural model quality. Briefings in Bioinformatics, 25(2): 387–395, 2024. doi: 10.1093/bib/bbae064.

15. J.Michael Word, Simon C. Lovell, Thomas H. LaBean, Hope C. Taylor, Michael E. Zalis, Brent K. Presley, Jane S. Richardson, and David C. Richardson. Visualizing and quantifying molecular goodness-of-fit: small-probe contact dots with explicit hydrogen atoms11edited by j. thornton. Journal of Molecular Biology, 285(4): 1711–1733, 1999. ISSN 0022-2836. doi: 10.1006/jmbi.1998.2400.

16. Jens Wiedemann, Tobias Zok, Marko Milostan, et al. LCS-TA to identify similar fragments in RNA 3D structures. BMC Bioinformatics, 18:456, 2017.

17. Pengyu Xiong, Rui Wu, Jian Zhan, et al. Pairing a high-resolution statistical potential with a nucleobase-centric sampling algorithm for improving RNA model refinement. Nature Communications, 12(1): 2777, 2021. doi: 10.1038/s41467-021-23100-4.

18. S. Zhang, Y. Liu, and L. Xie. A universal framework for accurate and efficient geometric deep learning of molecular systems. Scientific Reports, 13:19171, 2023. doi: 10.1038/s41598-023-46382-8.

19. Jun Li, Wei Zhu, Jun Wang, Wenfei Li, Sheng Gong, Jian Zhang, and Wei Wang. RNA3DCNN: Local and global quality assessments of RNA 3D structures using 3D deep convolutional neural networks. PLOS Computational Biology, 14(11):1–18, 11 2018. doi: 10.1371/journal.pcbi.1006514.

20. Clément Bernard, Guillaume Postic, Sahar Ghannay, and Fariza Tahi. RNA-TorsionBERT: leveraging language models for RNA 3D torsion angles prediction. Bioinformatics, 41(1), January 2025. doi: 10.1093/bioinformatics/btaf004.

21. Sandro Bottaro, Francesco Di Palma, and Giovanni Bussi. The Role of Nucleobase Interactions in RNA Structure and Dynamics. Nucleic acids research, 42, 10 2014. doi: 10.1093/nar/gku972.

22. Adam Zemla, Ceslovas Venclovas, John Moult, and Krzysztof Fidelis. Processing and analysis of CASP3 protein structure predictions. Proteins: Structure, Function, and Bioinformatics, 37(S3):22–29, 1999. doi: 10.1002/(SICI)1097-0134(1999)37:3+<22::AID-PROT5>3.0.CO;2-W.

23. Tomasz Zok, Mariusz Popenda, and Marta Szachniuk. MCQ4Structures to compute similarity of molecule structures. Central European Journal of Operations Research, 22, 04 2013. doi: 10.1007/s10100-013-0296-5.

24. M. Biasini, T. Schmidt, S. Bienert, V. Mariani, G. Studer, J. Haas, N. Johner, A. D. Schenk, A. Philippsen, and T. Schwede. Openstructure: an integrated software framework for computational structural biology. Acta Crystallographica Section D: Biological Crystallography, 69(5): 701–709, 2013. doi: 10.1107/S0907444913007051.

25. C. Zhang, M. Shine, A. M. Pyle, and Y. Zhang. US-align: Universal Structure Alignment of Proteins, Nucleic Acids and Macromolecular Complexes. Nature Methods, 19: 1109–1115, 2022.

26. J. Abramson, J. Adler, J. Dunger, et al. Accurate structure prediction of biomolecular interactions with AlphaFold 3. Nature, 2024. doi: 10.1038/s41586-024-07487-w.

27. Yusuke Kagaya, Zhen Zhang, Nur Muhammad Ibtehaz, et al. NuFold: end-to-end approach for RNA tertiary structure prediction with flexible nucleobase center representation. Nature Communications, 16:881, 2025. doi: 10.1038/s41467-025-56261-7.

28. Tianqi Shen, Zhenpeng Hu, Sheng Sun, et al. Accurate RNA 3D structure prediction using a language model-based deep learning approach. Nature Methods, 21: 2287–2298, 2024. doi: 10.1038/s41592-024-02487-0.

29. Y. Li, C. Zhang, and C. Feng. Integrating end-to-end learning with deep geometrical potentials for ab initio RNA structure prediction. Nature Communications, 14:5745, 2023. doi: 10.1038/s41467-023-41303-9.

30. Emidio Capriotti, Tomas Norambuena, Marc A. Marti-Renom, and Francisco Melo. All-atom knowledge-based potential for RNA structure prediction and assessment. Bioinformatics, 27(8):1086–1093, 02 2011. ISSN 1367-4803. doi: 10.1093/bioinformatics/btr093.

31. Dirk Merkel. Docker: lightweight linux containers for consistent development and deployment. Linux journal, 2014(239): 2, 2014.

32. R. C. Kretsch, E. S. Andersen, J. M. Bujnicki, W. Chiu, R. Das, B. Luo, B. Masquida, E. K. S. McRae, G. M. Schroeder, Z. Su, J. E. Wedekind, L. Xu, K. Zhang, I. N. Zheludev, J. Moult, and A. Kryshtafovych. RNA target highlights in CASP15: Evaluation of predicted models by structure providers. Proteins, 91(12):1600–1615, Dec 2023. doi: 10.1002/prot.26550.

33. Bohdan Schneider, Blake Alexander Sweeney, Alex Bateman, et al. When will RNA get its AlphaFold moment? Nucleic Acids Research, 51(18): 9522–9532, 2023. doi: 10.1093/nar/gkad726.

34. Jeremy Wohlwend, Gabriele Corso, Saro Passaro, Noah Getz, Mateo Reveiz, Ken Leidal, Wojtek Swiderski, Liam Atkinson, Tally Portnoi, Itamar Chinn, Jacob Silterra, Tommi Jaakkola, and Regina Barzilay. Boltz-1: Democratizing Biomolecular Interaction Modeling. 2025. bioRxiv doi: 10.1101/2024.11.19.624167, 26 April 2025, pre-print: not peer-reviewed.

35. Clément Bernard, Guillaume Postic, Sahar Ghannay, and Fariza Tahi. Has AlphaFold 3 achieved success for RNA? Acta Crystallographica Section D, 81(2), 2025. doi: 10.1107/S2059798325000592.

