## Supplementary file for "RNAdvisor 2: A unified platform for RNA 3D model quality assessment using metrics, scoring functions, and meta-metrics"

#### RNA quality assessment

##### A. Metrics

***RNA-oriented metrics.***

**Longest Continuous Segments in Torsion Angle space (LCS-TA)**

The longest continuous segments in torsion angle space (LCS-TA) is a metric that is based on the MCQ metric. It computes the longest continuous segments that have similar torsional angle values given a threshold (between 10, 15, 20 or 25 degrees). As the metric is performed on torsional space, it does not require any superimposition. This metric is based on a divide and conquer procedure, where a continuous window is sliding over the torsional angle space. It starts with the length of the sequence, before dividing the window into two equal parts. The process is repeated until the window size can not be divided.

##### B. Scoring functions

***Statistical potentials.***

**RNA-BRiQ**

RNA-BRiQ (Backbone Rotameric and Quantum-mechanical-energy-scaled base-base knowledge-based potential) is a fully knowledge-based scoring function primarily used for robust RNA structure refinement. It adopts a nucleobase-centric approach, treating each base as a rigid body. The core concept of RNA-BRiQ is capturing the complete orientation dependence of key interactions. It models interactions between bases (base-base), between a base and a main-chain oxygen atom (base-oxygen), and between two main-chain oxygen atoms (oxygen-oxygen). These orientation-dependent potentials are derived from structural statistics collected from known RNA structures. A significant novelty is the use of quantum mechanical (QM) calculations to reweight the base-base statistical potentials, aiming to minimize the effects of indirect interactions present in empirical data. The backbone is modelled using rotameric states and empirical internal energies. The total energy $E$ is the sum of six terms:

$$E = E_{\text{bb}} + E_{\text{bo}} + E_{\text{oo}} + E_{\text{rot}} + E_{\text{internal}} + E_{\text{clash}}$$

with $E_{\text{bb}}$, $E_{\text{bo}}$ and $E_{\text{oo}}$ potentials that capture orientation dependence using kernel density estimations derived from known structures. The $E_{\text{bb}}$ potentials are scaled by QM calculations performed on representative base pair configurations extracted from PDB structures. The $E_{\text{rot}}$ models backbone using statistical rotameric states for ribose and dihedral angle coupling for phosphate. $E_{\text{internal}}$ accounts for bond lengths, bond angles, and dihedral angles between the phosphate group and connecting riboses. $E_{\text{clash}}$ is an atomic clash energy based on distances and empirical parameters. The base-base orientation is described in a six-dimensional space using distance vectors and rotational angles. RNA-BRiQ's potentials were derived from all high-resolution RNA structures from the PDB (resolution <3.0Å) to maximize the available data for collecting base and backbone statistics. The key advantages of RNA-BRiQ include its ability to refine RNA structures, the incorporation of quantum mechanical to capture orientation dependence, and the use of a nucleobase-centric sampling algorithm. It is, like other knowledge-based methods, limited by the quality and diversity of the training dataset.

***Deep learning.***

**LociPARSE**

lociPARSE (1) is a deep learning model inspired by AlphaFold 2 (2) that predicts the LDDT score of RNA 3D structures. lociPARSE consists of multiple IPA (Invariant Point Attention, an attention mechanism that applies to 3D geometry) layers that process the input RNA 3D structure to refine nucleotide and pair features. Following the IPA layers, a linear layer and a two-layer fully connected network (MLP) are used to estimate nucleotide-wise Local Distance Difference Test (LDDT) scores, which are then aggregated to predict the global structural accuracy. The model was trained on a dataset derived from trRosettaRNA (3), containing 1,399 RNA targets and 51,763 structural models generated by various RNA 3D structure prediction methods. It can output a confidence score for each nucleotide in the predicted structure, indicating the local accuracy of the predicted structure. lociPARSE uses several input features, including the one-hot encoding of the nucleotide, its relative position in the sequence, the sequential separation of nucleotide pairs (discretized into bins representing different interaction ranges), and the interatomic distances between all pairs of $P$, $C_4'$, and glycosidic $N$ atoms, encoded using Gaussian radial basis functions. These features are designed to be invariant to global Euclidean transformations.

**PAMNet**

The Physics-Aware Multiplex Graph Neural Network (PAMNet) (4) is a graph neural network (GNN) framework designed for accurate and efficient representation learning of 3D molecules across different sizes and types. Is is not specifically designed for RNA but can be applied to any 3D molecule. PAMNet represents each molecule as a two-layer multiplex graph, distinguishing between local and non-local interactions based on principles from molecular mechanics. It uses global and local message passing modules to update node embeddings, incorporating geometric information such as pairwise distances and angles. A fusion module with a two-step attention pooling process combines the updated embeddings for downstream tasks. The input features for PAMNet vary depending on the task, but it uses atomic numbers (Z) as initial node features and encodes pairwise distances and angles using basis functions.

**TB-MCQ**

The TorsionBERT-MCQ (5) is a method based on a deep learning method that leverages language models to predict the torsion angles of RNA structures. It has been derived from the RNA-TorsionBERT that predicts RNA torsional angles from the sequence. Given a structure, we extract the torsional angles and use the predicted torsional angles to then compute the MCQ (6) between the predicted and the native structure. The model has been trained on a dataset of 4267 structures derived from the PDB. This scoring function only considers the torsional angles and not the distance between atoms, which might be limited if used alone to assess the quality of a structure.

### Benchmark

#### C. Link between metrics

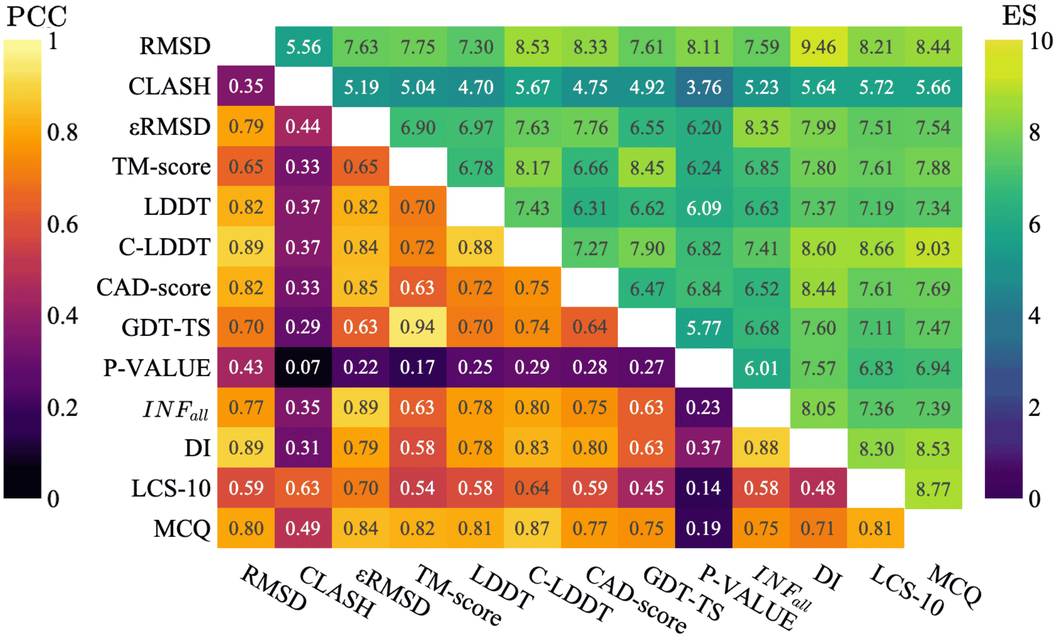


**Figure S1.** ES and PCC scores for each metric on Test Set I. The lower half of the matrix represents the PCC, while the upper half corresponds to the ES score. The diagonal has a PCC of 1 and ES of 10.

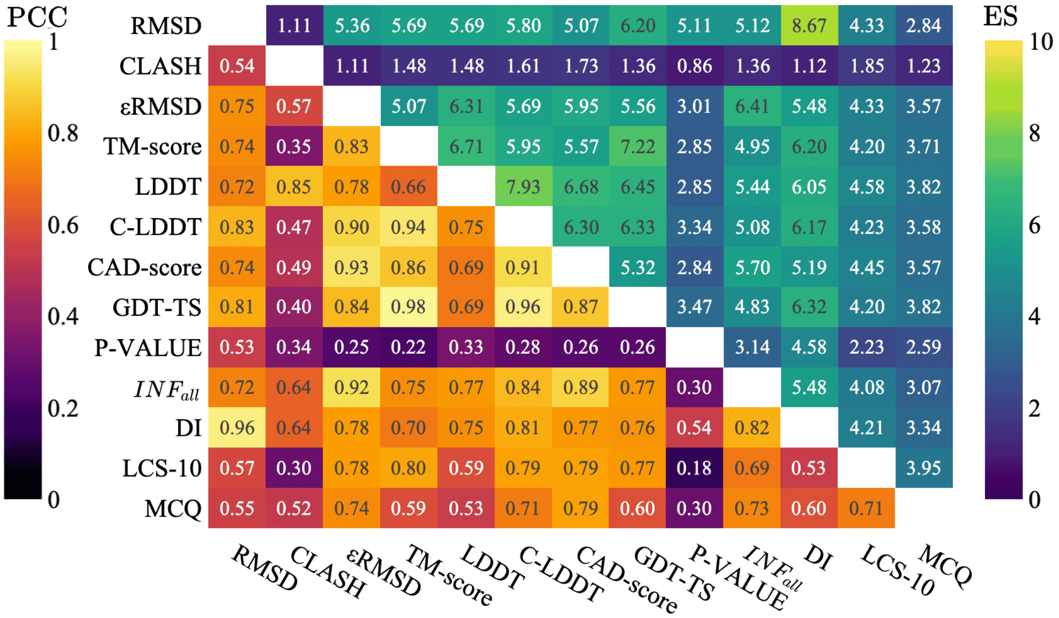


**Figure S2.** ES and PCC scores for each metric on Test Set II. The lower half of the matrix represents the PCC, while the upper half corresponds to the ES score. The diagonal has a PCC of 1 and ES of 10.

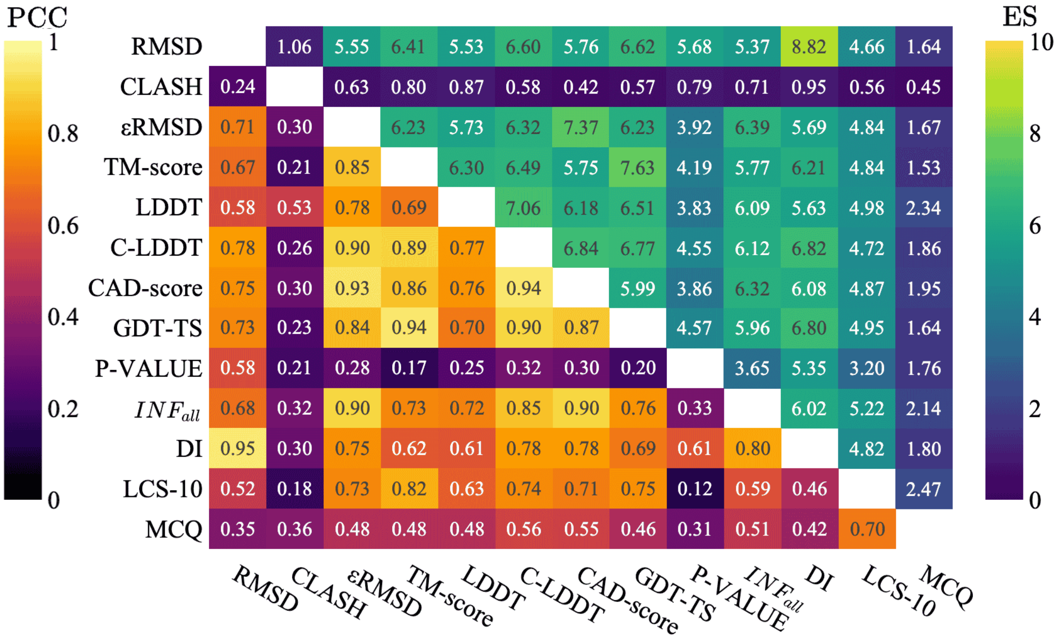


**Figure S3.** ES and PCC scores for each metric on Test Set III. The lower half of the matrix represents the PCC, while the upper half corresponds to the ES score. The diagonal has a PCC of 1 and ES of 10.

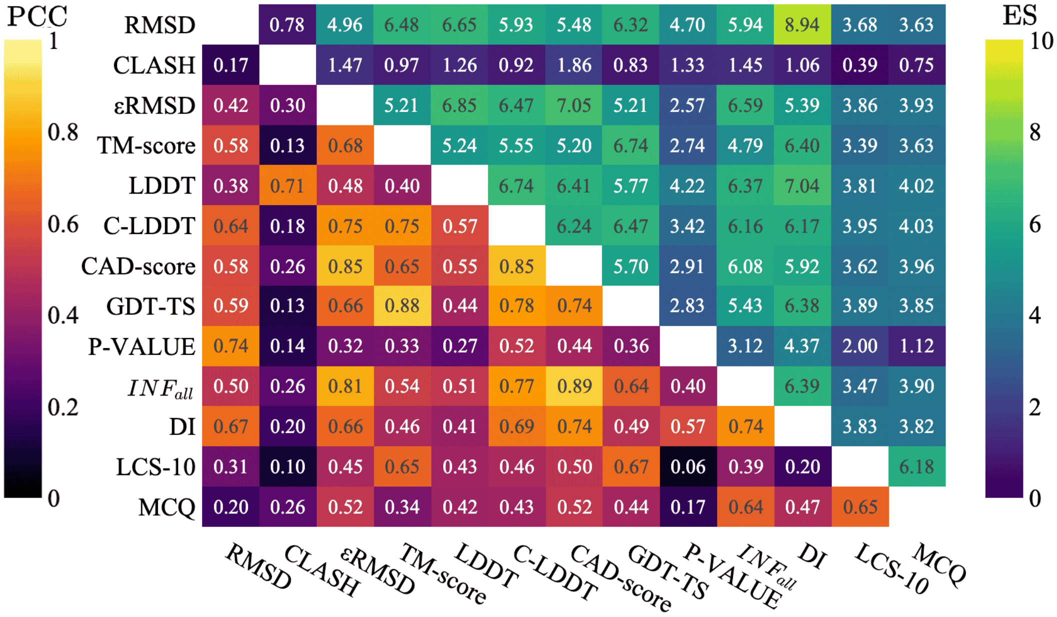


**Figure S4.** ES and PCC scores for each metric on Test Set IV. The lower half of the matrix represents the PCC, while the upper half corresponds to the ES score. The diagonal has a PCC of 1 and ES of 10.

#### D. Meta-metrics and metrics

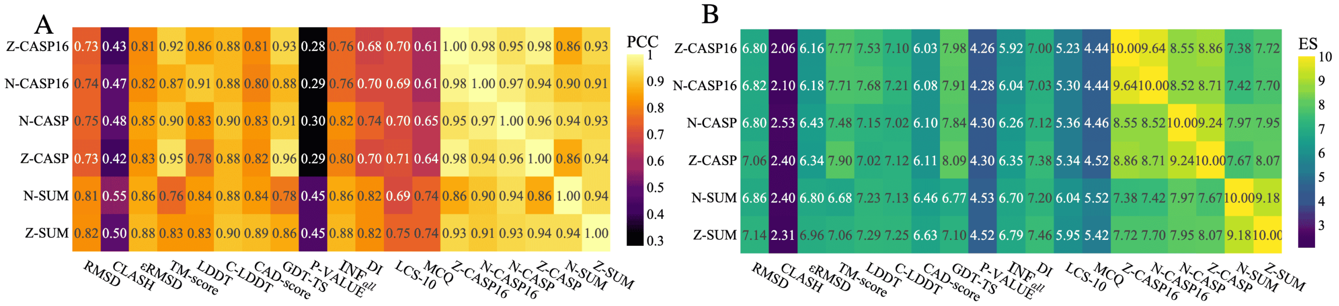

**Figure S5.** Comparison of six meta-metrics (Z-SUM, N-SUM, Z-CASP, N-CASP, Z-CASP16, N-CASP16) with existing metrics in terms of A) PCC and B) ES. Values are averaged over the four test sets.

The large number of available evaluation metrics can make it difficult to compare and rank predicted structures consistently. To simplify this process, we introduce meta-metrics that combines multiple metrics into a single, interpretable score. We compared the six meta-metrics with the existing metrics to see if they are correlated and if they can be used to assess the quality of a structure.

The results are shown in Figure S5, with the PCC shown in Figure S5A and the ES in Figure S5B. The six meta-metrics considered are the Z-SUM, a Z-score computed on the sum of all eleven metrics (we omitted the C-LDDT as it is redundant with the LDDT); N-SUM, its min-max normalised counterpart, Z-CASP, the CASP15 weighted Z-score, N-CASP which applies the same CASP weights but with min-max normalisation, Z-CASP16, the CASP16 weighted Z-score, and N-CASP16 with the min-max normalisation. The results both demonstrate that the meta-metrics are generally well correlated with individual metrics, confirming their capacity to capture consistent structural quality. In particular, Z-SUM exhibits high correlations across nearly all metrics (PCC higher than 0.8 for most, except MCQ and LCS-10), and achieves a high enrichment score (between 6 and 7), indicating it is especially effective at selecting top-performing structures across diverse quality measurements. The correlation with the CLASH-score remains very low (PCC around 0.5 and ES around 2), which makes sense as CLASH-score is less correlated to the other metrics and has a weight of $\frac{1}{11}$ in the overall sum. Furthermore, the meta-metrics are strongly intercorrelated (PCC higher than 0.9). This supports the idea that these aggregations consistently reflect structural quality despite different normalisation strategies or weightings. Results for each dataset are provided in Figures S6, S7, S8 and S9 of the Supplementary file.

Based on its overall performance in correlation and enrichment analyses, we adopt the Z-SUM as the reference meta-metric for RNA 3D structure quality assessment in the remainder of this study. We also include the N-SUM for further comparative analyses when we want to have a more interpretable metric.

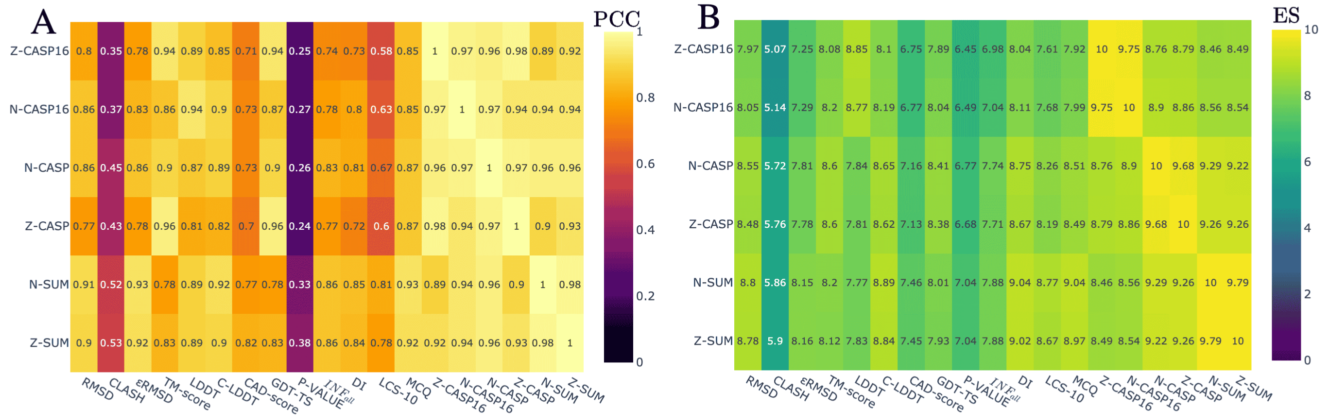

**Figure S6.** Comparison of six meta-metrics (Z-SUM, N-SUM, Z-CASP, N-CASP, Z-CASP16, N-CASP16) with existing metrics in terms of A) PCC and B) ES for Test Set I.

#### Scoring functions and metrics relationship

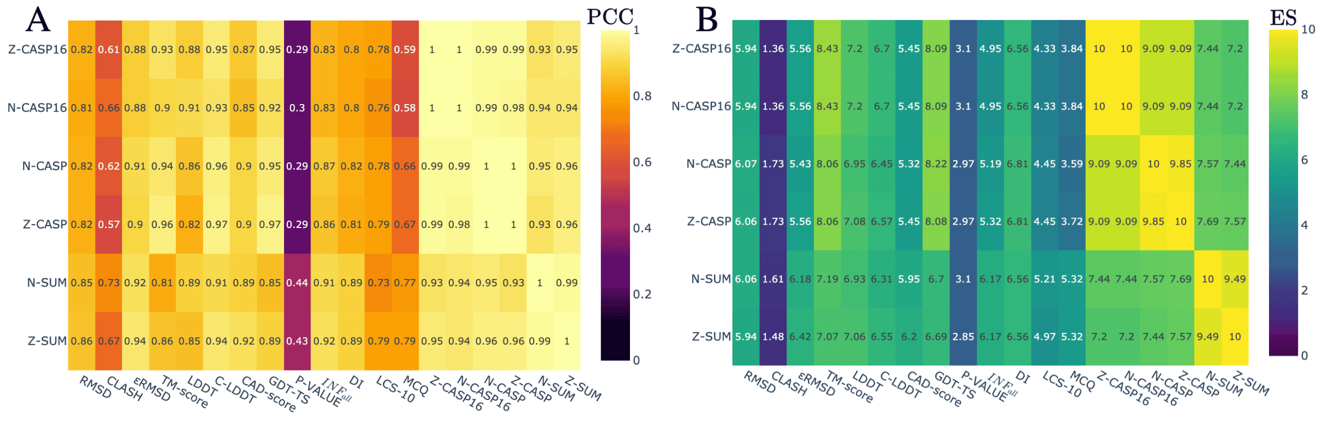


**Figure S7.** Comparison of six meta-metrics (Z-SUM, N-SUM, Z-CASP, N-CASP, Z-CASP16, N-CASP16) with existing metrics in terms of A) PCC and B) ES for Test Set II.

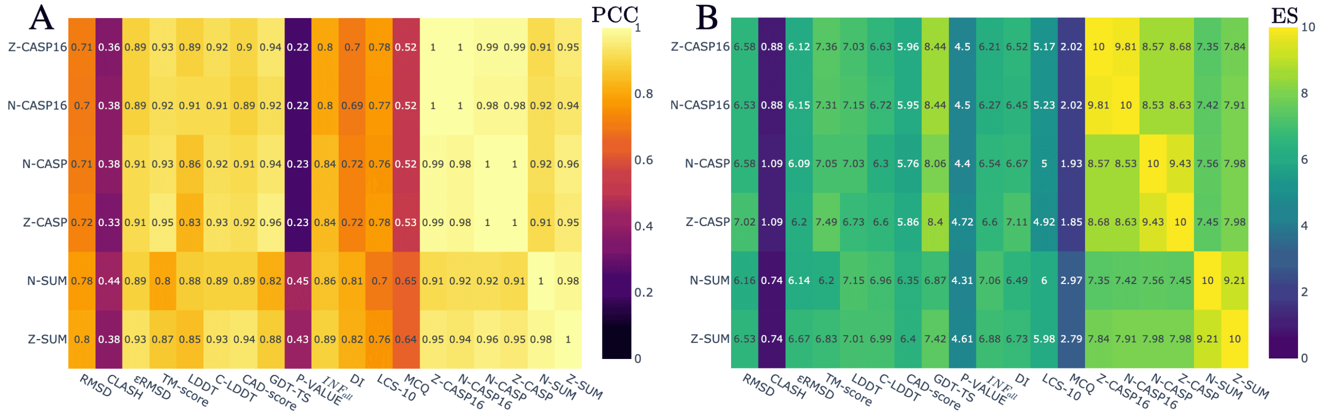


**Figure S8.** Comparison of six meta-metrics (Z-SUM, N-SUM, Z-CASP, N-CASP, Z-CASP16, N-CASP16) with existing metrics in terms of A) PCC and B) ES for Test Set III.

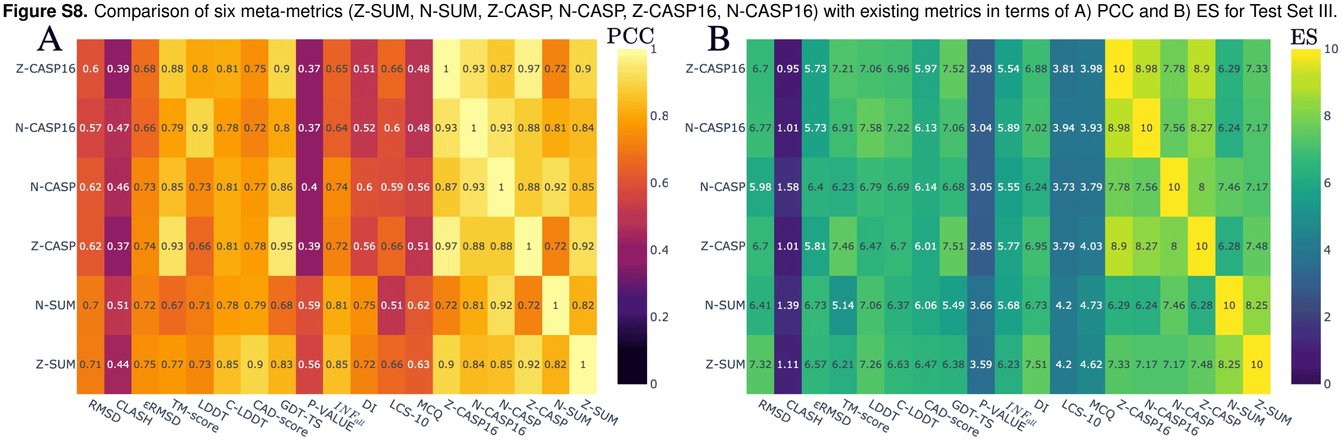


**Figure S9.** Comparison of six meta-metrics (Z-SUM, N-SUM, Z-CASP, N-CASP, Z-CASP16, N-CASP16) with existing metrics in terms of A) PCC and B) ES for Test Set IV.

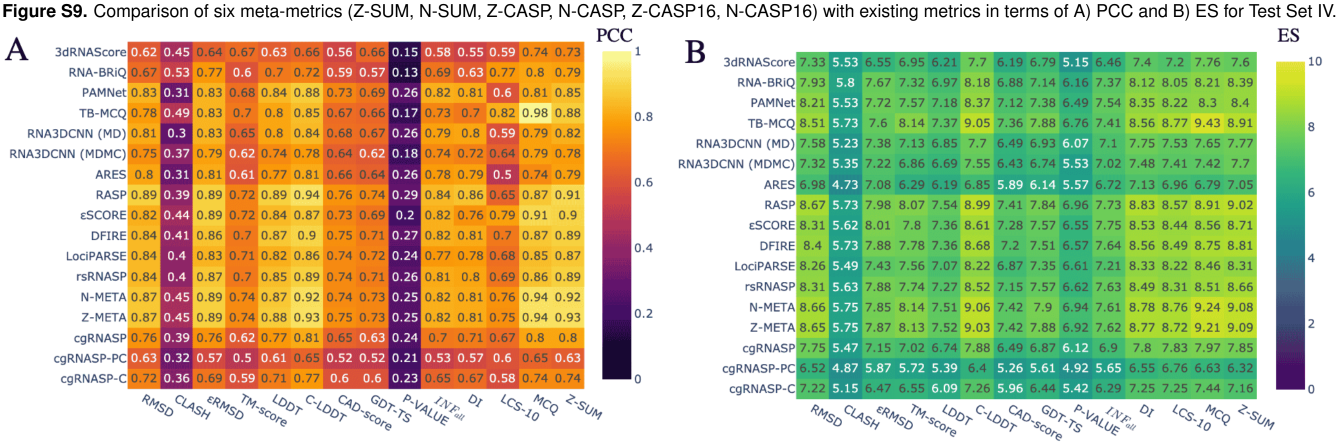


**Figure S10.** Link between the scoring functions and the metrics on Test Set I. A) PCC and B) ES scores. Scoring functions are sorted by the Z-SUM (last column).

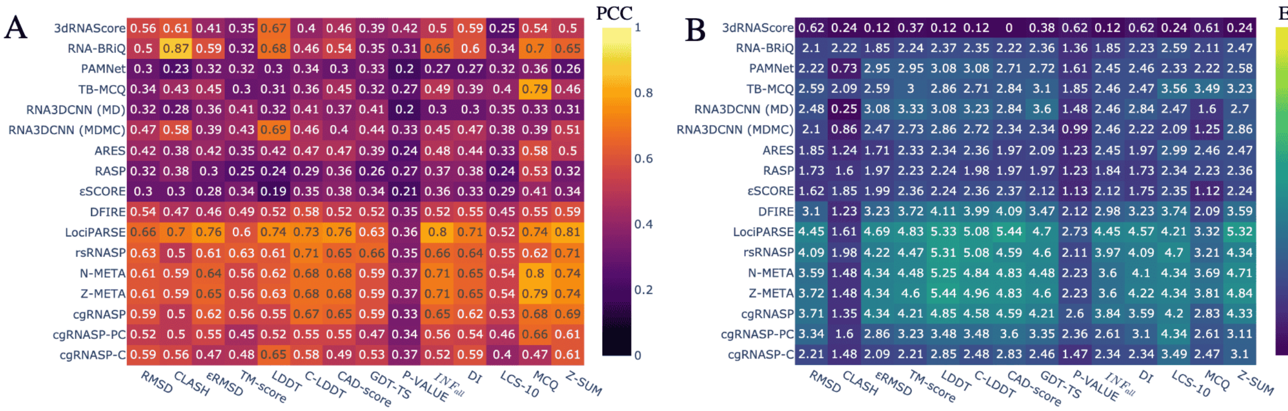


**Figure S11.** Link between the scoring functions and the metrics on Test Set II. A) PCC and B) ES scores. Scoring functions are sorted by the Z-SUM (last column).

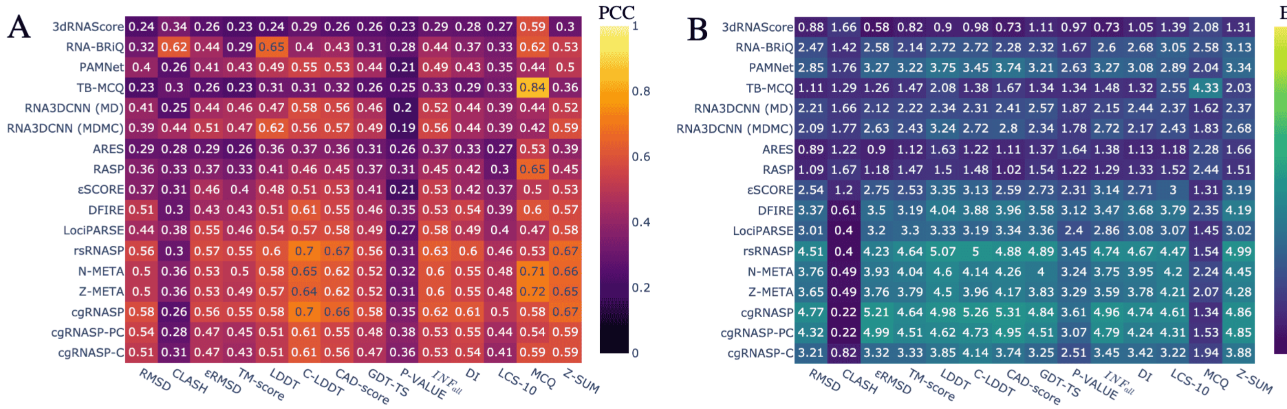


**Figure S12.** Link between the scoring functions and the metrics on Test Set III. A) PCC and B) ES scores. Scoring functions are sorted by the Z-SUM (last column).

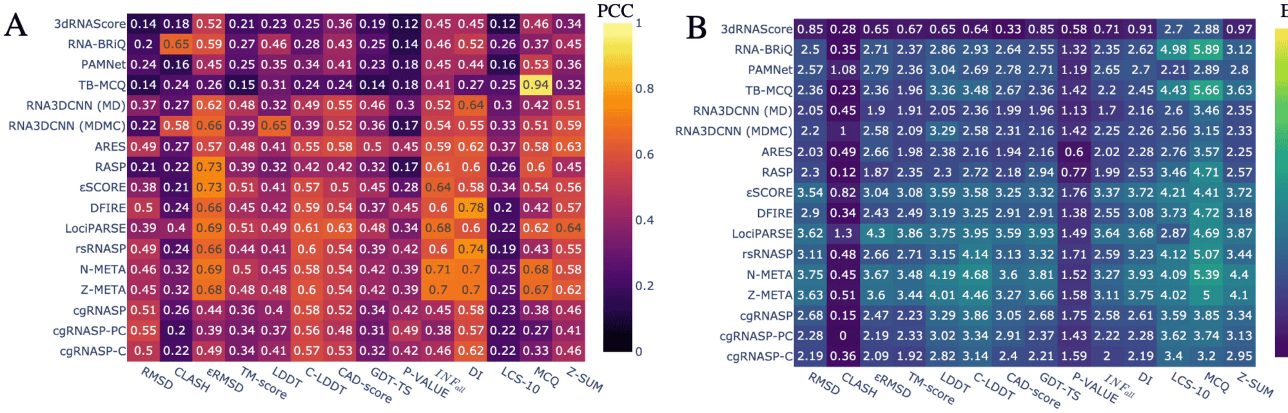


**Figure S13.** Link between the scoring functions and the metrics on Test Set IV. A) PCC and B) ES scores. Scoring functions are sorted by the Z-SUM (last column).

**Table S1.** Mean N-SUM of the best structure based on each scoring function for each Test set. The maximum of N-SUM is 11 (corresponds to the normalised sum of eleven metrics). The methods are ranked by the overall mean of N-SUM over the four test sets.

| | TSI | TSII | TSIII | TSIV | Mean |
|---|---|---|---|---|---|
| rsRNASP (7) | 7.31 | **9.02** | 9.14 | 5.66 | **7.78** |
| cgRNASP (8) | 6.97 | 8.70 | **9.15** | 5.64 | 7.62 |
| Z-META | 7.26 | 8.44 | 7.75 | 5.45 | 7.22 |
| LociPARSE (1) | 6.72 | 8.91 | 6.72 | 6.27 | 7.15 |
| N-META | 7.16 | 8.15 | 7.75 | 5.45 | 7.13 |
| cgRNASP-PC (8) | 6.24 | 7.97 | 8.42 | 5.40 | 7.01 |
| RNA-BRiQ (9) | 7.30 | 7.08 | 7.07 | **6.41** | 6.96 |
| DFIRE (10) | 7.25 | 7.67 | 7.40 | 5.04 | 6.84 |
| $\epsilon$SCORE (11) | 7.34 | 7.04 | 6.10 | 6.02 | 6.62 |
| PAMNet (4) | 7.28 | 6.72 | 6.25 | 5.82 | 6.52 |
| RNA3DCNN (MDMC) (12) | 6.64 | 5.14 | 7.79 | 5.52 | 6.27 |
| TB-MCQ (5) | **7.42** | 6.56 | 5.63 | 5.31 | 6.23 |
| cgRNASP-C (8) | 6.11 | 6.83 | 7.05 | 4.51 | 6.13 |
| RASP (13) | 7.29 | 5.16 | 5.10 | 5.38 | 5.73 |
| RNA3DCNN (MD) | 6.79 | 6.17 | 6.32 | 3.32 | 5.65 |
| ARES (14) | 5.84 | 5.67 | 4.88 | 5.18 | 5.39 |
| 3dRNAScore (15) | 7.01 | 2.68 | 4.87 | 5.05 | 4.90 |

To observe the ranking quality for the different scoring functions, we can also consider the N-SUM of the best structure based on the scoring function ranking. We provide in Table S1 the N-SUM for the top structure ranked by each scoring function for each dataset. We use the N-SUM (compared to the Z-SUM) to easily interpret the results: the values range between 0 and 11, where 11 is the best score. For the Test Set I (with near-native decoys), the best method is TB-MCQ, with a mean N-SUM of 7.42, which is surprising as it is a torsional-based scoring function. As it did not find the native structure (28 out of 85), the top structure has a good score, but is not the native one. It might be explained by the decoys' creation, which will create near-native structures without good torsional angles compared to the other dataset decoys. rsRNASP has the highest mean N-SUM for Test Set II (9.02), and cgRNASP has the highest mean N-SUM for Test Set III (9.15), followed closely by rsRNASP (N-SUM of 9.14). For Test Set IV, the best scoring function is RNA-BRiQ with a mean N-SUM of 6.41, which remains lower than the other datasets. The best scoring functions based on the four test sets are rsRNASP, cgRNASP and Z-META (mean of N-SUM of 7.78, 7.62 and 7.22).

### Bibliography


1. S. Tarafder and D. Bhattacharya. lociparse: A locality-aware invariant point attention model for scoring rna 3d structures. *Journal of Chemical Information and Modeling*, 64(22):8655–8664, Nov 2024. doi: 10.1021/acs.jcim.4c01621.
2. John Jumper, Richard Evans, Alexander Pritzel, Tim Green, Michael Figurnov, Olaf Ronneberger, Kathryn Tunyasuvunakool, Russ Bates, Augustin Žídek, Anna Potapenko, Alex Bridgland, Clemens Meyer, Simon A. A. Kohl, Andrew J. Ballard, Andrew Cowie, Bernardino Romera-Paredes, Stanislav Nikolov, Rishub Jain, Jonas Adler, Trevor Back, Stig Petersen, David Reiman, Ellen Clancy, Michal Zielinski, Martin Steinegger, Michalina Pacholska, Tamas Berghammer, Sebastian Bodenstein, David Silver, Oriol Vinyals, Andrew W. Senior, Koray Kavukcuoglu, Pushmeet Kohli, and Demis Hassabis. Highly accurate protein structure prediction with AlphaFold. *Nature*, 596:583–589, 8 2021. ISSN 0028-0836. doi: 10.1038/s41586-021-03819-2.
3. W. Wang, C. Feng, and R. Han. trRosettaRNA: automated prediction of RNA 3D structure with transformer network. *Nat Commun*, 14:7266, 2023. doi: 10.1038/s41467-023-42528-4.
4. S. Zhang, Y. Liu, and L. Xie. A universal framework for accurate and efficient geometric deep learning of molecular systems. *Scientific Reports*, 13:19171, 2023. doi: 10.1038/s41598-023-46382-8.
5. Clément Bernard, Guillaume Postic, Sahar Ghannay, and Fariza Tahi. RNA-TorsionBERT: leveraging language models for RNA 3D torsion angles prediction. *Bioinformatics*, 41(1), January 2025. doi: 10.1093/bioinformatics/btaf004.
6. Tomasz Zok, Mariusz Popenda, and Marta Szachniuk. MCQ4Structures to compute similarity of molecule structures. *Central European Journal of Operations Research*, 22, 04 2013. doi: 10.1007/s10100-013-0296-5.
7. Ya-Lan Tan, Xunxun Wang, Ya-Zhou Shi, Wenbing Zhang, and Zhi-Jie Tan. rsRNASP: A residue-separation-based statistical potential for RNA 3D structure evaluation. *Biophysical Journal*, 121: 142–156, 1 2022. ISSN 00063495. doi: 10.1016/j.bpj.2021.11.016.
8. Ya-Lan Tan, Xunxun Wang, Shixiong Yu, Bengong Zhang, and Zhi-Jie Tan. cgrnasp: coarse-grained statistical potentials with residue separation for rna structure evaluation. *NAR Genomics and Bioinformatics*, 5(1), 03 2023. ISSN 2631-9268. doi: 10.1093/nargab/lqad016. lqad016.
9. Pengyu Xiong, Rui Wu, Jian Zhan, et al. Pairing a high-resolution statistical potential with a nucleobase-centric sampling algorithm for improving RNA model refinement. *Nature Communications*, 12(1):2777, 2021. doi: 10.1038/s41467-021-23100-4.
10. Emidio Capriotti, Tomas Norambuena, Marc A. Marti-Renom, and Francisco Melo. All-atom knowledge-based potential for RNA structure prediction and assessment. *Bioinformatics*, 27(8): 1086–1093, 02 2011. ISSN 1367-4803. doi: 10.1093/bioinformatics/btr093.
11. Sandro Bottaro, Francesco Di Palma, and Giovanni Bussi. The Role of Nucleobase Interactions in RNA Structure and Dynamics. *Nucleic acids research*, 42, 10 2014. doi: 10.1093/nar/gku972.
12. Jun Li, Wei Zhu, Jun Wang, Wenfei Li, Sheng Gong, Jian Zhang, and Wei Wang. RNA3DCNN: Local and global quality assessments of RNA 3D structures using 3D deep convolutional neural networks. *PLOS Computational Biology*, 14(11):1–18, 11 2018. doi: 10.1371/journal.pcbi.1006514.
13. Emidio Capriotti, Tomas Norambuena, Marc A. Marti-Renom, and Francisco Melo. All-atom knowledge-based potential for RNA structure prediction and assessment. *Bioinformatics*, 27:1086–1093, 4 2011. ISSN 1460-2059. doi: 10.1093/bioinformatics/btr093.
14. Raphael J. L. Townshend, Stephan Eismann, Andrew M. Watkins, Ramya Rangan, Maria Karelina, Rhiju Das, and Ron O. Dror. Geometric deep learning of RNA structure. *Science*, 373:1047–1051, 8 2021. ISSN 0036-8075. doi: 10.1126/science.abe5650.
15. Jian Wang, Yunjie Zhao, Chunyan Zhu, and Yi Xiao. 3dRNAscore: a distance and torsion angle dependent evaluation function of 3D RNA structures. *Nucleic Acids Research*, 43:e63–e63, 5 2015. ISSN 1362-4962. doi: 10.1093/nar/gkv141.
